# A Hierarchical Reinforcement Learning Model Explains Individual Differences in Attentional Set Shifting

**DOI:** 10.1101/2021.10.05.463165

**Authors:** Anahita Talwar, Quentin Huys, Francesca Cormack, Jonathan P Roiser

**Affiliations:** Neuroscience and Mental Health Group, UCL Institute of Cognitive Neuroscience, 17-19 Queen Square, London, WC1N 3AZ; Max Planck UCL Centre for Computational Psychiatry and Ageing Research, Russell Square House, London, WC1B 5EH; Cambridge Cognition Ltd, Tunbridge Court, Bottisham, Cambridge, CB25 9TU; Division of Psychiatry, University College London, Maple House, 149 Tottenham Court Rd, London W1T 7BN

## Abstract

Attentional set shifting refers to the ease with which the focus of attention is directed and switched. Cognitive tasks such as CANTAB IED reveal great variation in set shifting ability in the general population, with notable impairments in those with psychiatric diagnoses. The attentional and learning processes underlying this cognitive ability, and how they lead to the observed variation remain unknown. To directly test this, we used a modelling approach on two independent large-scale online general-population samples performing CANTAB IED and psychiatric symptom assessment. We found a hierarchical model that learnt both feature values and dimension attention best explained the data, and that compulsive symptoms were associated with slower learning and higher attentional bias to the first relevant stimulus dimension. This data showcase a new methodology to analyse data from the CANTAB IED task, and suggest a possible mechanistic explanation for the variation in set shifting performance, and its relationship to compulsive symptoms.

## Introduction

When faced with unfamiliar challenges, we may look to reapply previously successful strategies, or switch to exploring novel ones. The ease with which we can generalise pre-learnt action-outcome associations, and the flexibility with which we can grasp previously irrelevant ones depends on how and what we learn to attend to. These cognitive faculties known as attentional set formation, and attentional set shifting, respectively, have commonly been assessed with the CANTAB Intra-Extra Dimensional Set Shift Task (Owen et al., 1991). The task, originally designed as a computerised analogue of the Wisconsin Card Sort Task (Berg, 2010; Grant & Berg, 1948), consists of several stages requiring participants to use trial and error to learn which feature of a multidimensional stimulus signals the correct response for that stage of the task. The introduction of novel stimuli occurs twice, first to assess set formation by the reapplication of previously learned rules, and second to assess the crucial attentional set shift by switching attentional focus to the previously irrelevant stimulus dimension. This task has been used extensively to document cognitive impairments in patients with psychiatric diagnoses. Early research identified difficulties in performing attentional set shifts but not simpler reversals, as measured by increased errors, in patients such as those with frontal lobe excisions suggesting that attentional set shifts represent a distinct and complex higher-level process (Owen et al., 1991). Difficulties in set shifting have also been considered a cognitive hallmark of OCD (Chamberlain SR et al., 2006, 2007; Purcell R et al., 1998; Vaghi et al., 2017; Veale DM et al., 1996) exemplified by the use of CANTAB IED as an endpoint in clinical studies (Tyagi et al., 2019). Despite this, there have been conflicting reports regarding the nature of these difficulties (Gottwald et al., 2018). Furthermore, difficulties in attentional set shifting have been reported in depression (Purcell et al., 1998; Purcell R et al., 1997), schizophrenia (Elliott R et al., 1995; Levaux MN et al., 2007; Liang S et al., 2018) and anxiety (Kim KL et al., 2019), highlighting that these alterations in cognition are not specific to OCD patients. Here, we will consider the methodologies that may have contributed to the discrepancies and non-specificity previously reported, and explore the extent to which a computational model of CANTAB IED can resolve these.

In recent years, the vast heterogeneity within psychiatric diagnoses has been highlighted (Fried EI, 2017; Fried EI & Nesse RM, 2015) and put forward as an explanation for inconsistent results in casecontrol studies. This has motivated the idea that symptoms of common mental health disorders in the general population are best described as falling along continuous dimensions, rather than by subdivisions into distinct categories. This conceptualisation has stimulated the measurement of transdiagnostic self-reported symptoms in the context of academic research (Cuthbert & Insel, 2013). Only a few studies using CANTAB IED have taken this dimensional approach. For example, Laws and colleagues focused solely on delusion symptoms in healthy controls, finding that more delusion-prone individuals made more errors on the early reversal learning stages of IED (Laws et al., 2011). An attractive experimental method that aligns with the dimensional framework involves the collection of very large datasets online to leverage the variation inherent in large samples and assess the relationship between symptoms and cognitive measures (Gillan et al., 2016).

In addition to the aforementioned clinical heterogeneity, variability in cognitive task performance is also likely to be underpinned by mechanistic heterogeneity. Traditional coarse measures of overall task performance, such as errors on the extra-dimensional shift stage, do not take into account patterns of trial-by-trial choices or the complex attention and learning mechanisms that are thought to produce them, and are therefore limited in the mechanistic insight they can provide. The advent of computational modelling in cognition aims to enhance the understanding of cognitive task performance by building generative models that explain trial-by-trial choices (Adams et al., 2016; Montague et al., 2012). The multidimensional nature of CANTAB IED task stimuli means that we cannot easily interpret the reasons for participants stimulus choices, and therefore what leads to variations in set shifting performance. Building generative models can help to provide insights by describing the attentional biases to stimulus dimensions that might develop throughout the task, how these influence learning and ultimately whether these processes account for set shifting performance. Furthermore, as these models estimate the level of randomness in participants’ choices, they can provide more precise participant-specific measures in the form of a small number of interpretable model parameters. Though increasingly popular in mental health research, no computational models have been built to describe the full IED task yet.

The theory-driven field of computational psychiatry involves developing models by mathematically specifying our hypotheses of the cognitive processes involved in performing the task that best describe variation in symptoms between participants. Reinforcement learning models, in which agents use feedback to learn actions that maximise their total reward (Sutton & Barto, 1998), have been extensively applied to cognitive tasks where learning is involved, including those assessing cognitive flexibility (Daw et al., 2011b). For example, Niv and colleagues developed the “dimensions task” to explore how selective attention aids learning about complex stimuli (Niv et al., 2015). It is similar to CANTAB IED in that participants are shown multidimensional stimuli and have to use feedback to infer the ‘correct’ feature on each block of the task. The authors showed that models of reinforcement learning on stimulus features fit participants’ choices the best (Niv et al., 2015), and subsequently that attention biased both learning and decision making on this task (although attention was measured by eye-tracking and was not mathematically specified: (Leong et al., 2017)). Despite the similarities between the individual stages of these tasks, the different ordering of the stages means that the tasks are measuring distinct attentional process. In the dimensions task, the correct feature on each stage is randomly chosen, independent of other stages, whereas in IED, the ‘correct’ feature is from the same dimension for the first seven stages of the task, which strongly promotes the formation of an attentional set. Additionally, it has been suggested that attentional set shifts rely on transfer to novel exemplars, which do not truly exist in the dimensions task, meaning that intra- and extra-dimensional set shifts cannot be well-defined (Slamencka NJ, 1968). The clearer separation of these processes in IED allows for a more straightforward interpretation of attentional set formation and cognitive flexibility (Downes et al., 1989). Furthermore, the dimensions task has not been used extensively in clinical populations, highlighting the utility of developing models for the CANTAB task. Only one study using CANTAB IED has investigated reinforcement learning models, although they focused on the simple discrimination and reversal learning stages at the beginning of the task (Murray GK et al., 2008). The authors did not fit computational models to trial-by-trial data but used error scores from the relevant task stages to infer that schizophrenia patients exhibit an impairment in basic reinforcement learning. As the analysis did not include the crucial set shifting stages, we are unable to draw conclusions about the underlying processes affecting their performance from this study.

In this paper we present the first attempt to develop a full computational model of CANTAB IED that incorporates the set shifting stages and estimates participant-specific parameters by fitting to trial-by-trial data. We use a large dataset of unselected volunteers, tested online, and explore the mechanistic insights that the models provide. We validate the model comparison on a second large dataset of healthy volunteers and analyse how the model parameters relate to symptoms of common mental health disorders, focusing on OCD. Our modelling approach is inspired by previous models of the ‘dimensions task’ including feature-based reinforcement learning with attention mechanisms to explore whether these models are also able to capture participants’ choices on the crucial set shifting stages.

## Materials and Methods

### Participants

Two independent datasets were collected online via Prolific Academic by Cambridge Cognition Ltd. Participants were recruited if they confirmed they a) were over 18 years of age, b) were fluent in English, c) had not experienced a significant head injury (resulting in loss of consciousness), d) reported not having been diagnosed with an untreated mental health condition (by medication or psychological intervention) that had a significant impact on their daily life, e) had never been diagnosed with mild cognitive impairment or dementia. All participants were paid at a rate of £7.50 per hour. Participants’ data was anonymous and they provided their consent online before participating in the experiment. The first dataset includes 731 participants who completed the IED task. The second dataset includes 762 participants who completed the IED task and also several self-report mental health questionnaires. These sample sizes provide 95% power to detect associations of r = 0.13 at alpha = 0.05 (two-tailed).

### CANTAB Intra-Extra Dimensional Set Shift Task

To assess attentional set shifting, participants completed the CANTAB Intra-Extra Dimensional Set Shift Task (IED), originally designed as a computerised analogue of the Wisconsin Card Sort Task (Berg, 2010; Grant & Berg, 1948). The design of the task is presented in Figure 1. On each trial, participants were presented with a choice between two stimuli, which for most of the task are compound stimuli comprising two dimensions – lines and shapes. The chosen stimulus was indicated with a mouse click. Selecting one of the stimuli resulted in deterministic feedback indicating whether their choice was correct. Participants were instructed to use this feedback to learn an underlying rule. On achieving six correct choices in a row, it is assumed the rule has been learnt, and they move on to the next stage where there is a new rule. If a participant completed 50 trials on a stage without achieving six correct choices in a row, the stage is failed and the task terminated. Throughout the task, the rule is that one of the features indicates the correct stimulus. Critically, this feature is from the line dimension for Stages 1-7, and the shape dimension in Stages 8-9 (in the second dataset, this was reversed such that the correct feature was from the shape dimension for Stages 1-7, and from the line dimension for Stages 8-9). During the task, participants must learn reversals (same stimuli but reversed such that the other feature of the same dimension becomes correct), intra-dimensional shifts (new stimuli and the correct feature is from the same dimension) and extra-dimensional shifts (new stimuli and the correct feature is from the other dimension). An increase in errors on Stage 8 (extra-dimensional shift) is typical as participants find it difficult to attend to the previously irrelevant dimension. No participants were excluded for the analysis of task data.

**Figure 1.**
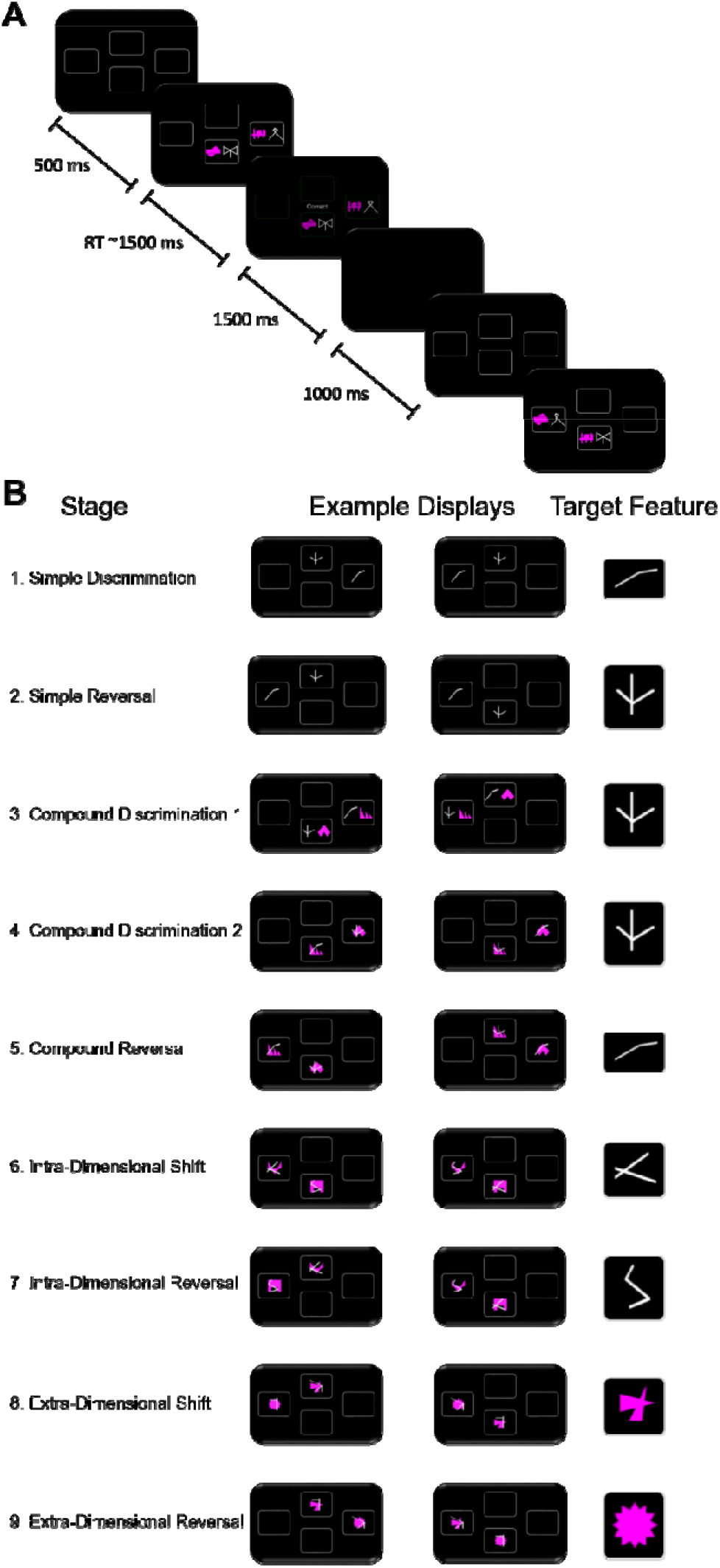
CANTAB IED Task Schematic. **A.** Schematic of single example trial from Stage 3 of CANTAB IED. Participants are presented with two stimuli that are composed of one feature from each of two dimensions: pink shapes, and white lines. Participants select a stimulus and receive deterministic feedback that informs them of whether their choice was correct or incorrect. After the feedback, the screen briefly turns blank before the presence of two new stimuli indicates the start of a new trial. **B.** Illustration of all nine stages of CANTAB IED, displaying two example trials from each stage, as well as the target features for each stage. Participants need to learn that this feature indicates the correct stimulus for each trial of that stage.

### Self-Report Questionnaires

In the second dataset, participants were additionally asked to provide their age, gender and level of education^1^. Participants also completed questionnaires assessing compulsive symptoms (Obsessive Compulsive Inventory Revised, (Foa et al., 2002)), depressive symptoms (Self-Rating Depression Scale,(Zung WWK, 1965)), anxious symptoms (State Trait Anxiety Inventory, (Spielberger et al., 1989), and schizotypy symptoms (Short Scales for Measuring Schizotypy,(Mason O et al., 2005)).

### Identifying clusters (K-means)

For each participant in the second dataset, an ‘error trajectory’ was determined as the number of errors at each of the stages of the task. K-means clustering using the *sklearn.cluster.KMeans* package in Python, version 3.7.1 was applied to these trajectories, treating the trajectory as a multidimensional point. This divides participants into a prespecified number (K) clusters, based on the trajectory of their errors over the course of the task. Each participant was allocated to the cluster with the nearest mean trajectory. The algorithm was run 10 times with different initial mean values, and the best fitting output from these runs was used as the final clustering. This clustering was also used to predict cluster labels of model-simulated data for each participant, given their best-fitting parameters. Only participants with both human and model-simulated data from all nine stages of the IED task could be included in this analysis, leaving 611 participants. K was chosen to be three based on the screeplot of the sum of squared distances of samples to their closest cluster centre. The participants excluded from the K-means analysis formed an additional cluster, giving four behaviourally defined clusters.

### Computational models

CANTAB IED data are traditionally analysed in terms of the number of errors per stage. However, these summary statistics do not make full use of the richness inherent in the dataset. Computational models, on the other hand, model the trial-by-trial choices of each participant, and therefore capture the underlying attention and learning dynamics that are necessary to complete the task. Thus, we developed computational models to capture relevant attentional and learning processes to directly test whether they account for the variation in set shifting behaviour. Initial models were based on reinforcement learning (see below), whilst subsequent models included an additional layer where weights represent the allocation of attention to different stimulus dimensions. Analysis of the parsimony of these models in explaining participant data was used to guide development of the next model. The three main models are described below.

#### 1. Feature Reinforcement Learning (fRL)

This model calculates the values (V) of stimuli (S) by summing the weights (W) of features (f) present in the stimuli:

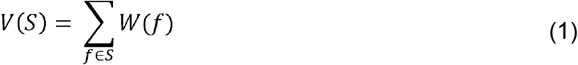

All feature weights are initialised to 0. The stimulus values are then entered into a softmax probabilistic choice function:

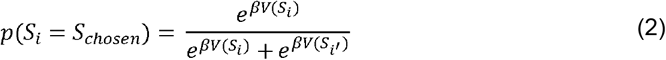

where β is the inverse temperature parameter, such that large β leads to more deterministic choices of the higher-valued stimulus, and small β leads to more random decisions that are less dependent on stimulus values. The model uses a reinforcement learning rule (Sutton and Barto, 1998) to update the weights of features in *both* stimuli on each trial, as follows:

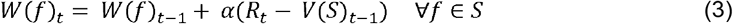

This model does not treat each stimulus as independent, as it takes into account that stimuli share features. However, it does not consider that the features are part of two different dimensions, and thus cannot generalise across dimensions. Therefore, the next model incorporated an attentional component to estimate stimulus values, allowing it to capture within-dimension generalisation to novel stimuli.

Free parameters: α (learning rate), β (choice determinism)

#### 2. Combined Attention-Modulated Feature Reinforcement Learning (Ca-fRL)

This model incorporates dimensional attentional weights to account for the attentional biases towards stimulus dimensions that might develop throughout the task, and capture the within-dimension generalisation to novel stimuli. These weights play a role in stimulus valuation, where they multiply feature weights from the corresponding dimension, thereby weighing the contribution of features from different dimensions according to how much attention is being paid to each one:

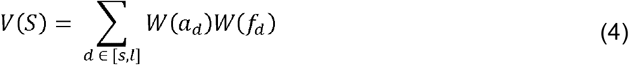

More precisely, the attention weight for the initially relevant dimension is specified as [σ(θ)], and for the other dimension as [1-σ(θ)], where σ indicates a transformation by the sigmoid function. Thus, if one of these weights differs from 0, the other will also, but in the other direction. When both weights are close to 0, the dimensions are weighed equally. For instance, in the first dataset the line dimension was the initially relevant one, and stimulus values were calculated as follows:

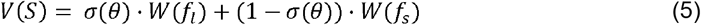

Stimulus values are then entered into a softmax probabilistic choice function as in the fRL model. All feature weights are initialised to 0. The initial value of θ (θ_0_) is a free parameter inferred from the data and represents a dimension primacy effect - the extent to which the initially relevant dimension is attended to, and the second dimension is ignored when introduced on Stage 3. Standard backpropagation updates the hidden weights and θ on each trial by differentiating the squared error loss (L) with respect to the weight being updated:

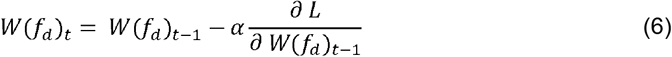

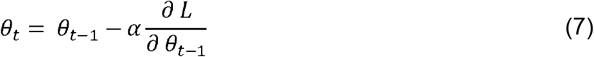

Backpropagation of the hidden weights involves multiplication by θ. Therefore, when θ is high (more attention to initially relevant dimension), the weights of features from the other dimension will be updated less on that trial, and vice versa; thus capturing the attentional processes missing from the fRL model. Notably, the same learning rate is used to update the feature and dimension weights, suggesting that a *combined* process underlies the learning taking place.

Free parameters: α (learning rate), β (choice determinism), θ_0_ (dimension primacy)

#### 3. Separate Attention-Modulated Feature Reinforcement Learning (Sa-fRL)

This model is identical to Ca-fRL except that a different learning rate, ε, is used to update θ:

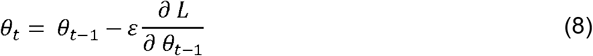

The hidden weights are updated using α, as in equation 6. This model allows us to test the possibility that identifiably *separate* learning rates underlie the learning of features values and dimensional attention allocation, which cannot be captured by the combined learning model.

Free parameters: α (learning rate – features), ε (learning rate – dimensions), β (inverse temperature), θ_0_ (dimension primacy)

### Parameter Estimation

We used a hierarchical Bayesian parameter estimation approach, described previously (Huys et al., 2011), which finds the maximum a posteriori parameter estimates for each participant, given the model and the data, and sets the parameters of the prior distribution to the maximum likelihood estimates given all participants’ data. The purpose of using hierarchical estimation here is mainly that priors over the parameters act to regularise the estimates such that unrealistic, extreme values are avoided. We used an expectation-maximisation approach which repeatedly iterates over two steps until convergence is reached. Briefly, on the E-step the model finds the best-fitting individual level parameter estimates for each participant given their data and the current parameters of the prior distribution, and on the M-step, the maximum likelihood group level prior parameters are updated to reflect the current individual parameter estimates.

All parameters were transformed before inference to ensure that 0 ≤ *α, ε* ≤ 1, and *β* ≥ 0. The mean and standard deviation for the best-fitting parameter values of each model are given in Table 1. Recoverability of parameters was calculated by simulating 500 datasets with parameters drawn randomly from the estimated prior distribution. The best-fitting parameters for these data sets were found as above, and parameter recoverability is indicated by the correlation between the simulated and recovered parameters (Table 1).

**Table 1.** Models + Parameter Recovery. Free parameters of each model, along with summary statistics of their best-fitting values and recoverability.

| Model | Parameter | Range | Mean (±SD) | Recovery |
| --- | --- | --- | --- | --- |
| fRL | Learning Rate | 0 - 1 | 0.62 ± 0.04 | 0.49 |
| | Choice Determinism | 0 - ∞ | 0.91 ± 0.33 | 0.90 |
| Ca-fRL | Learning Rate | 0 - 1 | 0.92 ± 0.14 | 0.84 |
| | Choice Determinism | 0 - ∞ | 1.34 ± 0.43 | 0.87 |
| | Dimension Primacy | $-\infty$ - ∞ | 1.78 ± 0.97 | 0.91 |
| Sa-fRL | Learning Rate - Features | 0 - 1 | 0.91 ± 0.14 | 0.88 |
| | Learning Rate - Dimensions | 0 - 1 | 0.85 ± 0.24 | 0.67 |
| | Choice Determinism | 0 - ∞ | 1.34 ± 0.42 | 0.87 |
| | Dimension Primacy | $-\infty$ - ∞ | 1.73 ± 0.87 | 0.91 |

**Table 2.** Relationship Between Parameters and Symptoms. Spearman’s rank correlations and p-values for relationship between symptom questionnaire scores and untransformed parameters of best-fitting models. p-values were calculated using a boostrapping procedure. OCI-R: Obsessive Compulsive Inventory-Revised; SDS: Symptoms of Depression Scale; STAI-S: State Trait Anxiety Inventory-State; STAI-T: State Trait Anxiety Inventory – Trait; SSMS: Short Scales for Measuring Schizotypy.

|  |  | Parameters |  |  |
| --- | --- | --- | --- | --- |
|  |  | Learning Rate | Choice Determinism | Dimension Primacy |
| Questionnaire | OCI-R | -.13, .0003 | -.09, .01 | .12, .0008 |
|  | SDS | -.04, .30 | -.04, .25 | .03, .42 |
|  | STAI-S | -.05, .18 | -.02, .55 | .05, .22 |
|  | STAI-T | -.01, .87 | .02, .63 | .00, .97 |
|  | SSMS | .01, .88 | .02, .50 | -.01, .77 |

### Model Comparison

A priori, we did not assume that any of the models should be more likely. Therefore, they can be compared by examining the approximate model log likelihood with the integrated Bayesian Information Criterion (Huys et al., 2011). This procedure gives us the model that fits the data most parsimoniously, whilst penalising for unnecessary added model complexity (additional parameters).

## Results

Participants completed CANTAB IED, a multidimensional two-alternative-forced choice task, in which one stimulus feature deterministically indicates the correct stimulus on each trial. Whilst the key feature changes on every stage of the task, it is from the same dimension for the first seven stages of the task, and switches to the other dimension on the crucial extra-dimensional set shift stage (Stage 8). Participants use trial and error to determine the underlying rule, and correct feature, for each stage of the task. Analysis of the number of errors made per stage on the CANTAB IED has been used extensively to identify differences in set shifting performance between diagnostic groups, however it is unable to account for the learning that takes place within stages, as well as the reasons behind stimulus choices (e.g., whether participants are focused on a particular dimension, or the whole stimulus). Here, we develop a modelling analysis that overcomes both limitations, and investigate how this relates to symptoms of common mental health disorders.

### Attention-Modulated Reinforcement Learning Accounts for Variation in ED Shift Performance

The raw human data in Figure 2. shows that the vast majority of participants make fewer than five errors on the first seven stages of the task. On Stage 8, when a feature from the previously irrelevant dimension is now completely predictive of the correct stimuli, there is a marked increase in variation in performance, with many participants failing this stage. We first tested a feature reinforcement learning model (fRL) that considers the dimensional composition of the task stimuli, which has been previously used to model data from similar tasks with multidimensional stimuli (Niv et al., 2015). This model learns weights for individual stimulus features and calculates the overall stimulus value by summing the weights of its component features, thereby accounting for the tendency to select a stimulus with a particular white line feature (for example), regardless of the pink shape (for example) that it is paired with, if participants believe that this feature currently indicates the correct stimulus.

**Figure 2.**
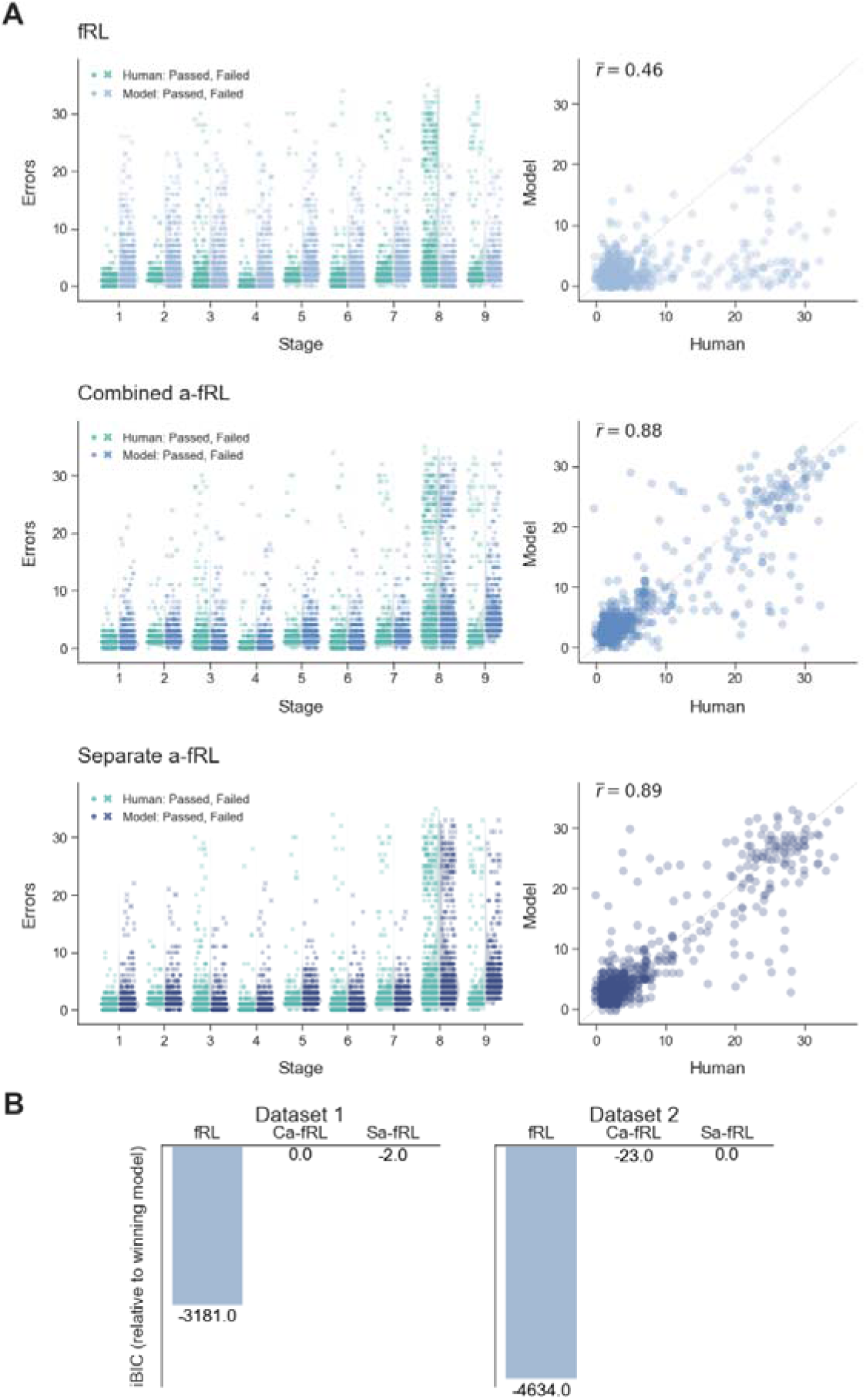
Qualitative and Quantitative Model Fits. **A.** Qualitative comparison of human dada with model simulated data for each participant given their best fitting parameters for models fRL (top), Ca-fRL (middle), and Sa-fRL (bottom). *Left:* Error distributions per stage of CANTAB IED. *Right:* ED Shift errors. Correlation coefficient indicates average between human data and 10 model simulated datasets. Jitter has been added to scatter points to aid visualisation. **B.** Quantitative model comparison with iBIC values for three models in two independent sets of data.

Figure 2A (top) shows fRL model simulated data and its differences from human data. The model-simulated error distributions match the human data fairly well for most stages, consistent with the notion that participants do not treat each stimulus as independent but take into account that stimuli share features. However, it is not able to capture the difficulty that many participants have on the ED Shift (Stage 8). Crucially, the model-simulated error distributions on the ID Shift (Stage 6) and ED Shift (Stage 8), in which participants are presented with novel stimulus features, are identical (and the same is true for their respective reversals: Stages 7 and 9). This is because the fRL model cannot take into account that the novel features are from the same dimensions, and thus does not generalise across dimensions, instead initiating new learning for unseen features. This pattern contrasts with the human data, where errors are generally lower when novel stimulus features are introduced but the relevant dimension remains the same (ID Shift), than when novel stimulus features are introduced and the relevant dimension changes (ED Shift). This is further evidenced by the relatively low correlation (r = 0.46) between human and model simulated errors on the ED Shift stage (Fig 2A top right). In summary, while the fRL model captures participants’ ability to weight stimulus features rather than whole stimuli, it does not capture their additional tendency to weight dimensions.

In order to account for dimension-based learning, we modified the above model to include a component that captures the dimension to which participants are attending. In line with previous work, this attention component biases both stimulus valuation, by overweighting the values of features from the more attended dimension, and learning, by updating the weights of features from the more attended dimension to a greater extent (Leong et al., 2017). The dimension attention weights themselves were updated using backpropagation. We tested two variations of this feature + attention reinforcement learning model, one with a single learning rate for both the feature weights and dimension weights (Ca-fRL), and one with a separate learning rate for each type of weight (Sa-fRL). Figure 2A (middle and bottom) shows that the addition of the attention layer markedly improves model performance, with qualitatively better matched distributions of errors between human and model-simulated data on the ID and ED Shift stages. This is confirmed by the greatly superior correlation between human and simulated datasets (Ca-fRL: r = 0.88, Sa-fRL: 0.89) on ED Shift errors.

Whilst the fits of Ca-fRL and Sa-fRL are hard to distinguish qualitatively using model simulations, a more formal model comparison considers both the model fit and the model complexity, providing an overall measure of model performance. The model comparison (Fig 2B) shows that Ca-fRL is the more parsimonious model in the first dataset, but that Sa-fRL is the more parsimonious model in the second dataset (indicated by the least negative iBIC score). As these iBIC values are very close, compared to the fRL model, it is not clear that the added dimension learning rate parameter in the Sa-fRL model sufficiently improves model fit to justify its added complexity. As we prefer simpler models for increased falsifiability and reduced overfitting, and due to the worse parameter recovery of the dimension learning rate parameter in the Sa-fRL model, we selected Ca-fRL as the most appropriate model for CANTAB IED data, thereby using it for our subsequent parameter analysis. Whilst it is somewhat surprising that such a simple learning model can capture almost all of the variation in set-shifting performance present in our sample, it is interesting that these kinds of models, which were initially created to describe how participants learn about multidimensional stimuli (Leong et al., 2017; Niv et al., 2015), are also able to account for set formation and shifting.

### Slower Learning, Random Choices and Stronger Dimension Primacy Lead to Difficulties in Set Shifting

In order to assess the Ca-fRL model’s predictions of overall task behaviour in more detail, we applied a K-means algorithm to cluster data based on participants’ errors-per-stage trajectories. The screeplot in Figure 3A shows that using more than three clusters does not reduce the sum of squared errors substantially, so a three-cluster solution was chosen. The largest cluster (cluster 1) is made up of participants that score few errors on all stages of the task. The other two clusters are made up of participants that made more errors on the ED shift, but are separated by their performance on the subsequent reversal. A final cluster (cluster 4) was added that was made up of the participants with incomplete IED task data (failed at Stage 8 or earlier) who could not be included in the K-means analysis. We then tested whether simulated data from each participant using the Ca-fRL model would fall into the same clusters as participants’ real data. Figure 3B shows that for the majority of participants, simulated data from their best fitting parameters using model Ca-fRL, was allocated to the same cluster as their original data. Some misalignment of real and model-simulated clustering is to be expected given that the K-means algorithm provides sharp cluster assignments, whereas real world behaviour cannot be so cleanly divided, and the assignment of some participants data is more ambiguous. The largest misalignment is of participants in cluster 3 to cluster 4, which might be because the small number of participants in cluster 3 (score high errors on the extra-dimensional shift and the subsequent reversal) passed the extra-dimensional set shift by chance rather than having obtained an understanding of the key rule change. Nonetheless, this approach provides further evidence that the model is able to capture the main features of and differences between participants’ task performance in our sample.

**Figure 3.**
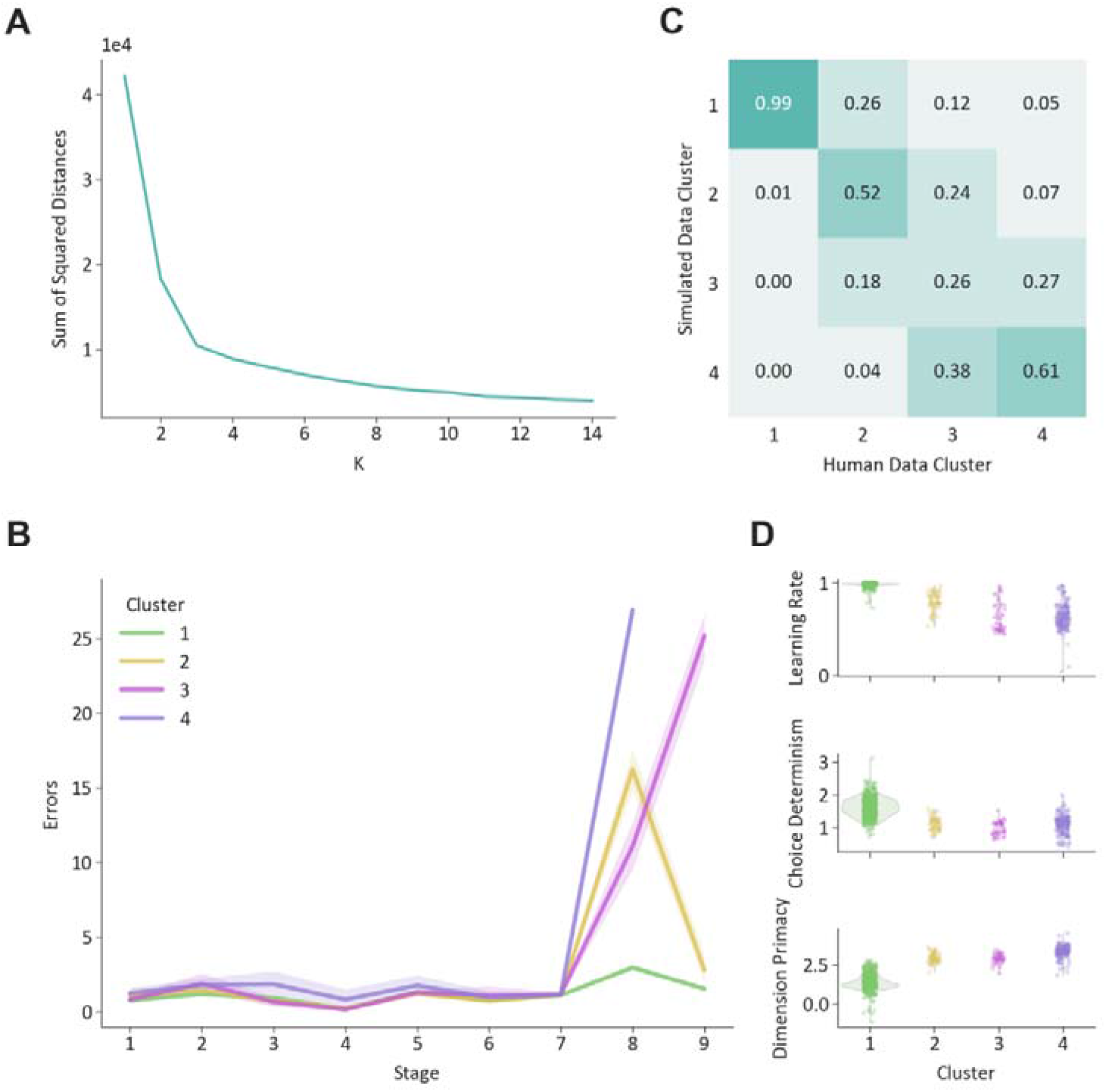
K-means Clustering of IED Data. **A.** K-means clustering of error per stage trajectories. Screeplot of the sum of square distances to cluster mean with K-means for different values of K. **B.** The clusters identified by a 3-cluster solution. The fourth cluster consists of participants who failed the task before reaching stage 9 (typically at stage 8, the ED shift) and could not be included in the analysis. Cluster sizes: Cluster 1: 532, Cluster 2: 50, Cluster 3: 42, Cluster 4: 138. **C.** Agreement between clustering of participants real and simulated data. Each column shows the proportions of simulated data (from participants in a particular human defined cluster) that were assigned to each cluster. **D.** Parameter distributions by clusters. Distributions of best fitting parameter values from Ca-fRL model, separated by cluster allocation.

To highlight the mechanistic insights that a modelling approach provides, we examined how model parameters varied by these behavioural clusters. Figure 3C shows that compared to the participants in cluster 1 (who score few errors throughout), participants in clusters 2, 3 and 4 have lower learning rates, choose more randomly, and pay more attention to the initially relevant dimension in the early stages of the task. More specifically, learning rates for participants in cluster 3 seem to be particularly low, whilst dimension primacy seems to be particularly high for participants in cluster 4, explaining why many in the latter cluster fail the extra-dimensional shift stage altogether.

### Slower Learning and Stronger Dimension Primacy Are Associated with Higher Compulsive Symptoms

To assess whether parameters from our best-fitting model, Ca-fRL, may be relevant to symptoms of mental illness, we examined their associations with a variety of symptom questionnaires: compulsivity, depression, state and trait anxiety, and schizotypy. Total score on the OCI-R questionnaire, which assess compulsive symptoms, was significantly associated with the learning rate parameter (r = -.13, p = .0003) and the dimension primacy parameter (r = .12, p = .0008) after applying Bonferroni correction (p < .003). Despite the high association between symptoms of mental health disorders, no other questionnaire scores were significantly correlated with model parameters. The specificity of these relationships was further highlighted with a Steiger’s Z test, which compares them to the next biggest correlation between parameters and symptoms scores, accounting for the association between the two symptom questionnaires of interest. For both parameters, this involved the relationship with state anxiety, and in both cases the association with compulsivity was significantly greater (learning rate: Z = 2.36, p = .009 and dimension primacy: Z = 2.26, p = .01).

To test whether these relationships were affected by the age, gender, or education level of participants, we conducted multiple regression analyses, predicting compulsivity symptoms from model parameters, including these variables as covariates. Again, learning rate and dimension primacy were significant predictors of OCI-R score when evaluated in separate models (learning rate: β = −9.85, 95% CI = [−14.30, −5.40], p = .00002, overall model F(4,757) = 16.14, R^2^ = 0.079; dimension primacy: β = 1.66, 95% CI = [0.86, 2.45], p = .00005, overall model F(4,757) = 15.54, R^2^ = 0.076). This confirms that participants who exhibit more compulsive symptoms tend to be slower to learn the task structure and show a greater attentional bias towards the initially correct stimulus dimension.

## Discussion

We implemented the first algorithmic analysis of the CANTAB Intra-Extra Dimensional Set Shift Task. We showed that a hierarchical reinforcement learning model with two simple levels provides a parsimonious yet highly accurate account for participant’s choices in two independent samples. In the model, lower-level weights represent the learnt values for stimulus features, and higher-level weights represent the learnt attention to stimulus dimensions. We also explored how model parameters were related to symptoms of common mental health disorders finding that lower learning rates were specifically associated with higher compulsive symptoms.

Our modelling analysis suggests a mechanistic explanation for how attention influences learning and decision making as well as for how the focus of attention is itself learnt and shifted. Our best-fitting model learns feature weights that represent current estimates of the associative value of features, and dimension attention weights that bias both the contribution of feature weights to stimulus valuation for action selection, but also the learning of the values of feature weights, as has been shown previously. Notably, our model extends previous algorithmic descriptions by suggesting that dimensional attention is itself learnt by simple prediction error-based update rules. More precisely, the dimension attention weights are updated based on tracking the predictive accuracy of the learnt values of their corresponding feature weights over time, in line with the idea that attention is directed to rewarding features (Mackintosh, 1975). Additionally, the rate of attentional learning is influenced by the strength of dimension attention itself, with more biased dimension attention slowing learning, in line with the idea that attention is updated faster when uncertainty is higher (Pearce & Hall, 1980). Thus, our results suggest that the attention is directed by both the expected reward and the uncertainty of stimuli lending support to hybrid models of attention (le Pelley et al., 2012). More specifically, our best-fitting model uses backpropagation to achieve this - updating feature and dimension weights to reduce error on each trial. Despite the superior performance of backpropagation-based algorithms for predicting human performance in a range of tasks (Kell AJE et al., 2018; Wenliang LK & Seitz AR, 2018), we acknowledge the historical scepticism around its implementation in the brain due to biological constraints (Crick, 1989). Recent research has offered biologically plausible approximations, such as using activity differences between sets of neurons in a local circuit to compute backprop-like weight updates using only locally available signals, providing new insights into possible implementations of backpropagation-like algorithms in the brain (Lillicrap et al., 2020).

The model suggests that a single type of underlying process, albeit in multiple instantiations, can explain performance across all IED task stages, including the extra-dimensional set shift. Reports that people with frontal lobe lesions, Parkinson’s Disease or obsessive compulsive disorder demonstrate impaired performance on the extra-but not intra-dimensional set shift or simple reversal learning stages (Downes et al., 1989; Owen et al., 1991; Purcell R et al., 1998) has led to the common assumption that the generalisation of pre-learnt rules to novel stimuli (intra-dimensional shift), and the shifting of attention or behaviour to new rules (extra-dimensional shift) are distinct cognitive processes. However, our model contains only a basic type of learning algorithm – albeit arranged hierarchically – and is able to account for choices on all task stages with multidimensional stimuli. There are two aspects to this finding. First, the need for a hierarchy does suggest the existence of separate processes. However, second, the fact that those processes are so similar suggests that the extra-dimensional set shifting process shares important attributes with simpler learning and reversal processes lower down in the hierarchy.

Some caveats to our results merit comment. First, the CANTAB IED has only one extra-dimensional set shift stage. This could result in noisier subject-specific parameter estimates and reduced sensitivity to identify more complex learning process, such as a separate learning rate parameter for the dimension weights. Typically modelling analysis of cognitive tasks involves repeated measurements of the cognitive construct of interest to obtain more reliable behavioural and algorithmic measurements. However, the introduction of novel stimuli and the specific ordering of task stages are thought to be crucial for the measurement of true attention set shifting and were intentionally considered in designing the task (Owen et al., 1991). Modelling analysis of tasks such as the ‘dimensions task’, akin to Wisconsin Card Sort, which do not involve the introduction of truly novel stimuli lends support to the idea of attention-weighted learning and decision-making. The use of our specific algorithm to update attention weights on these task versions is yet to be tested, and will be important to determine whether we could see evidence of a more complex process in a task with multiple set shifts. Second, our algorithm predicts higher errors on stage 9, the extra-dimensional reversal, compared to human data. This indicates that our model does not fully capture the flexibility with which humans make the extra-dimensional set shift. Relative inflexibility is a well-known feature of model-free reinforcement learning algorithms, which can be overcome by using model-based reinforcement learning algorithms or Kalman filter type models, suggesting an avenue for future research.

Analysis of associations with mental health questionnaire data revealed a specific negative relationship between the learning rate parameter and compulsivity. This suggests that participants with higher compulsive symptoms require more information to update their estimates and adapt their behaviour in light of changes in the environment. The majority of our sample had very high learning rates (between 0.95 and 1), which indicates that updates of value estimates after a single trial with unexpected feedback are sufficient to change behaviour. Such one-shot learning is indeed optimal in a deterministic environment. It is tempting to speculate that the ability to adapt behaviour swiftly relies on a better understanding of the structure of the task, or an impairment in participants with compulsive symptoms. These would be compatible with other modelling analyses of the probabilistic “two-step” task (Daw et al., 2011), which measures the extent to which participants arbitrate between ‘model-free’ and ‘model-based’ reinforcement learning systems. Whilst the former is relatively computationally cheap, it is slower due to reliance on learning from experienced feedback. The latter system benefits from fast learning and increased flexibility at the expense of increased computational complexity required to use a model or internal representation of the environment. A consistent finding is an overreliance of model-free behaviour in compulsive disorders (Gillan et al., 2016; Gillan CM et al., 2020; Voon et al., 2014).

Our sample consisted of unselected healthy volunteers. Despite the utility of online data collection, particularly for collecting a large number of participants, it is reliant on the pool of people who participate in online studies, which has the potential to introduce biases. Firstly, despite a very large sample relative to most studies in the field, the majority of participants performed very similarly, scoring few errors throughout the IED task. Similarly, the distributions of questionnaire scores were skewed towards the lower end of the scales, limiting the size of the relationship that we could detect in this sample. However, it is worth noting that the correlation between model parameters and compulsive symptoms is greater than that between ED Shift errors and compulsive symptoms, providing some evidence for the increased validity that a modelling approach is able to provide. Future research should focus on replicating this analysis in a sample with greater variation of symptom profiles.

In conclusion, we have shown that modelling analyses of CANTAB IED task performance are able to provide more precise explanations of behavioural differences. These explanations can then be leveraged to offer mechanistic insights into the symptoms of mental health disorders.

## Acknowledgements

The authors thank the MRC and Cambridge Cognition Ltd. who funded this work jointly through an MRC iCASE studentship grant MR/R015759/1. Cambridge Cognition employ FC and completed the data collection in this research.

## Footnotes

1 Left formal education before age 16, 2: Left formal education at age 16, 3: Left formal education at age 17-18, 4: Undergraduate degree or equivalent, 5: Master’s degree or equivalent, 6: PhD or equivalent.

